# Reducing uncertainty in impact assessments for alien species

**DOI:** 10.1101/2020.05.05.077958

**Authors:** David A. Clarke, David J. Palmer, Chris McGrannachan, Treena I. Burgess, Steven L. Chown, Rohan H. Clarke, Sabrina Kumschick, Lori Lach, Andrew M. Leibhold, Helen E. Roy, Manu E. Saunders, David K. Yeates, Myron P. Zalucki, Melodie A. McGeoch

## Abstract

Impact assessment is a widely used and cost-effective tool for prioritising invasive alien species. With the number of alien and invasive alien species expected to increase, reliance on impact assessment tools for the identification of species that pose the greatest threats will continue to grow. Given the importance of such assessments for management and resource allocation, it is critical to understand the uncertainty involved and what effect this may have on the outcome. Using an uncertainty typology and insects as a model taxon, we identified and classified the sources and types of uncertainty when performing impact assessments on alien species. We assessed 100 alien insect species across two rounds of assessments with each species independently assessed by two assessors. Agreement between assessors was relatively low for all three EICAT components (mechanism, severity, confidence) after the first round. For the second round, we revised guidelines and gave assessors access to each other’s assessments which improved agreement by between 20-30%. Of the 12 potential reasons for assessment discrepancies identified *a priori*, 11 occurred. The most frequent sources (and *types*) of uncertainty (i.e. differences between assessment outcomes for the same species) were: incomplete information searches (*systematic error*), unclear mechanism and/or extent of impact (*subjective judgment due to a lack of knowledge*), and limitations of the assessment framework (*context dependence*). In response to these findings, we identify actions to reduce uncertainty in the impact assessment process, particularly for assessing speciose taxa with diverse life histories such as Insecta. Evidence of environmental impact was available for most insect species, and (of the non-random original subset of species assessed) 14 or 29% of those with evidence were identified as high impact species (with either ‘Major’ or ‘Massive’ impact). Although uncertainty in risk assessment, including impact assessments, can never be eliminated, identifying and communicating its source and variety is a first step toward its reduction and a more reliable assessment outcome, regardless of the taxa being assessed.

## INTRODUCTION

With a changing climate and growing international trade, the range expansion of alien and invasive alien species is predicted to continue (Seebens et al. 2017). Ongoing arrival and establishment of alien species necessitates a triage approach to their management, where the most damaging, or those most likely to cause damage, are allocated high priority for surveillance and control (McGeoch et al. 2016). To this end, research on the impacts of alien species has been steadily growing (Crystal-Ornelas and Lockwood 2020), accompanied by the development of various assessment frameworks for classifying their impacts to assist in risk assessments (Roy et al. 2018, González-Moreno et al. 2019, Vilà et al. 2019).

Risk assessments are generally performed using multiple data types that vary in reliability, and therefore acknowledging, accounting and communicating associated uncertainty is an essential component of the process (Harwood and Stokes 2003). For example, assessments by the Intergovernmental Panel on Climate Change (IPCC), that use multiple lines of evidence, are conducted using a formal and agreed upon treatment of uncertainty (Mastrandrea et al. 2011). Impact assessments of alien species likewise should be accompanied by estimates of uncertainty (Roy et al. 2018), although this does not always occur (Caton et al. 2018). The environmental impact classification for alien taxa (EICAT), for example, requires assessors to provide a confidence rating (low, medium, high) for each piece of evidence used in the impact classification of each species (Hawkins et al. 2015). However, few studies have yet explored the potential types and sources of uncertainty in undertaking alien species impact assessments and how these may affect the final classification outcome (Kumschick et al. 2017a) who compared results from two different impact classification frameworks for alien amphibians). Better understanding of the sources of uncertainty in risk assessment is essential to protocol improvement, enabling more effective risk communication (McGeoch et al. 2012, Milner-Gulland and Shea 2017), and identifying general solutions for improving their reliability and value (Rueda-Cediel et al. 2018, Latombe et al. 2019).

While uncertainty cannot be removed entirely from risk assessment, identifying which sources of uncertainty are reducible is an essential step in improving the reliability and value of the information they generate. Given the inherent nature of uncertainty in ecology more broadly, and the need to account for it, frameworks that guide identifying and classifying various types of uncertainty have been found to be useful (Milner-Gulland and Shea 2017). For example, the uncertainty typology developed by Regan et al. (2002) has been used to assess uncertainty in conservation decision making in the face of climate change (Kujala et al. 2013), and to assess uncertainty in the alien species listing process and what effect it may have on estimating the identity and number of such species in countries (McGeoch et al. 2012). Regan et al. (2002) typology identified two broad types of uncertainty: epistemic and linguistic (for definitions see Methods), with multiple categories under each. Incorporation of both types covers the uncertainty in determinate facts, as well as that caused by the inherent variation in our use of language (Regan et al. 2002, Burgman 2005, Latombe et al. 2019). In invasion biology, language usage and terminology have been a significant hindrance to progress, including in the effective application of alien species risk and impact assessments (Roy et al. 2018, Vilà et al. 2019). Given the similarity between impact assessment procedures for alien species and components of ecological and invasion-focussed risk assessment (Yemshanov et al. 2012a, Yemshanov et al. 2012b), Regan et al. (2002) typology should likewise be suited to evaluating uncertainty in the assessment of alien species impacts.

Several alien invasive insect species pose significant risks to the environment and economy and require impact risk assessments to prioritize management (Bradshaw et al. 2016, Lovett et al.). Insects are one of the most species-rich, abundant, functionally important and yet understudied taxonomic groups (Stork et al. 2015, Noriega et al. 2018). This knowledge deficit extends to invasion ecology even though many alien insects are also some of the most economically important, in terms of both costs and benefits (Vilà et al. 2010, Bradshaw et al. 2016). Indeed, some alien species have become damaging after human introduction as managed species, e.g. *Bombus* species for managed pollination (Aizen et al. 2019). Despite a focus on impacts on agriculture, forestry and other commodities, a continually growing body of evidence demonstrates the importance of the detrimental effects of invasive alien insects on biodiversity and ecosystem functioning (Kenis et al. 2009, Herms and McCullough 2014, Brockerhoff and Liebhold 2017, Lester and Beggs 2019). Invasive ants, for example, are particularly well-known for outcompeting and displacing not only native ants, but having multiple indirect consequences (Holway et al. 2002), causing behavioural change in other taxa (Davis et al. 2008, Dorrestein et al. 2019), and significant declines in native species populations (O’Dowd et al. 2003, O’Connor et al. 2010). Other taxa, such as the hemlock woolly adelgid (*Adelges tsugae*) are responsible for largescale changes to native habitats (Spaulding and Rieske 2010). Continued appreciation of the potential and realised impacts of alien taxa has led to numerous government control programs targeting invasive arthropods, a substantial portion of which target insects (Suckling et al. 2019). Impact risk assessment for alien insects is therefore a large, challenging and important task, because of the numbers of species involved, gaps in knowledge of their life histories and distributions, and their substantial negative impacts. As a result, the delivery of robust and transparent impact assessments to support decisions about where to invest in and best prioritise intervention efforts is critical to successfully reducing the negative impacts of invasive alien species.

Here we identify and classify the types of uncertainty and their sources that may occur when assessing the environmental impacts of alien species. We do so by applying an existing assessment method (EICAT) using insects as the model taxon – a taxonomic group to which this method has not previously been applied. We evaluate the consequences of types of uncertainty on the level of congruence between independent assessments of the same species for 100 insect species. Incorporating the lessons learnt from performing EICAT assessments on alien insects, we conclude with recommendations for how best to communicate and mitigate certain types of assessment uncertainty.

## METHODS

For the purpose of identifying types of uncertainty in impact assessments for alien species, we used the semi-quantitative protocol, i.e. Environmental impact classification for alien taxa (EICAT). EICAT was developed to enable the mechanisms and severity of environmental impacts from alien taxa to be classified in a transparent, repeatable and comparative way (Blackburn et al. 2014, Hawkins et al. 2015), and has now been adopted by the IUCN as a standard for this purpose (IUCN). EICAT focuses exclusively on negative environmental impacts, specifically the most severe realised impacts, of alien species (not the overall net consequence that could include any positive effects) in their introduced range. The EICAT ‘severity of negative impacts’ are classified as ‘minimal concern’ (MC), ‘minor’ (MN), ‘moderate’ (MO), ‘major’ (MR), or ‘massive’ (MV) (Hawkins et al. 2015). The final severity attributed to an alien species at the end of the assessment process is its maximum realised impact anywhere within its introduced range, i.e. EICAT is only concerned with the most severe realised impact for a species. Two other categories are possible; ‘data deficient’ (DD) for cases where there is insufficient evidence to perform an assessment, or ‘not alien’ (NA) for when a given species does not in fact have an alien distribution.

EICAT assessments have, to date, been published for alien birds (Evans et al. 2016), amphibians (Kumschick et al. 2017a), selected gastropods (Kesner and Kumschick 2018) and mammals (Hagen and Kumschick 2018), and bamboo species (Canavan et al. 2019). Here we used the EICAT guidelines as published (Hawkins et al. 2015), although these guidelines have subsequently been revised by the IUCN via an online consultation process (IUCN). Our interest here was to identify forms of uncertainty in the impact assessment process, rather than to evaluate EICAT per se which is similar to a range of other impact assessment protocols (Kumschick et al. 2017b). We adopted previously suggested modifications to the mechanisms used to assess the environmental impacts of alien insects. Specifically, (i) the mechanism ‘Grazing/herbivory/browsing’ was altered to ‘Herbivory’ as grazing and browsing are not relevant for insects, and (González-Moreno et al. 2019) the addition of a relevant mechanism, ‘facilitation of native species’, was considered important for capturing this frequently encountered form of indirect impact of invasive insects on biodiversity (McGeoch et al. 2015).

The subset of alien insect species assessed was determined in the following way. With global alien insect species richness unknown, the initial universal set (∼2800 species) consisted of all insect species present in the Global Register for Introduced and Invasive Species (GRIIS), that provides verified species by country occurrences outside their native range and local evidence of impact ((Pagad et al. 2018), as at June 2017). In addition, alien insects known to impact the environment were extracted from relevant reviews (Kenis et al. 2009, Vaes-Petignat and Nentwig 2014, McGeoch et al. 2015, Cameron et al. 2016, Bertelsmeier et al. 2017, Evans et al. 2017). Species were then ranked by the number of times they were referred to across all sources, i.e. multiple sources designated them as ‘invasive’ or having a negative environmental impact. One hundred species were selected for assessment, including those qualitatively assessed as having most evidence of impact. Given the assumed positive relationship between pest severity and available literature on the species, we consider this suite of 100 species to incorporate the best-case scenario (Crystal-Ornelas and Lockwood 2020) for evidence-based assessment.

Eight of the 11 assessors were formally trained entomologists with significant relevant experience, and three were invasion biologists with substantial research experience in risk assessment (one specifically on EICAT) (co-authors on this paper). The qualifications and experience of the team of assessors was therefore comparable to EICAT conducted on other taxonomic groups to date (Evans et al. 2016, Kumschick et al. 2017b). In advance of the assessment exercise, the prior knowledge of each assessor of all 100 species in the subset was qualitatively self-assessed. Using a score of 0 – 5, where 0 is having never heard of a species and 5 is being very familiar with it, assessors gave a score for each species, followed by a confidence rating (high, medium, low). This exercise enabled an informed allocation of species to assessors, and species were thus assigned to assessors to as far as possible align most strongly with their specific expertise.

To measure the level of congruence between assessment outcomes and to identify reasons for the differences that occurred, each of the 100 insect species were each independently assessed by two assessors. The entire process can be generalised by the following steps. First, assessors discuss the method and perform a series of pilot assessments as a group to familiarise everyone with the EICAT process and protocol guidelines (Hawkins et al. 2015). Second, impact assessments are performed independently by two assessors for each species. Each species assessment results in three main outputs: 1) the mechanisms of impact (i.e. how the alien species is having a negative effect e.g. via competition), 2) the size of the impact (the EICAT ‘magnitude of impact severity’ category), and 3) the confidence in the evidence supporting the chosen mechanism and impact severity (Hawkins et al.)). Third, a revised and elaborated set of guidelines (Hawkins et al.)) was developed for assessing species based on the outcome of the first round where differences in outcomes occurred between assessors. Our intention here was to apply EICAT critically, adhering to the guidelines as far as possible and elaborating on or departing from these only where it was necessary to clarify, elaborate or modify details for the purpose of reducing uncertainty. The purpose of revisions to the guidelines was to clarify particular protocol points (Hawkins et al.)) that were identified as potentially resulting in differences between assessments, particularly to reduce the more common and readily addressed misunderstandings that arose in the first-round assessments (such as first confirming evidence of the existence of alien populations). Fourth, the revised guidelines, along with the initial results from both assessors for species where there were differences in the result of the assessment, are sent back to assessors. Differences can occur for one or more of the assessment outcomes per species, i.e. differences in allocated impact mechanism, severity, confidence level, or any combination of the three. Fifth, assessors then could either refine their initial assessment considering this new information, or to leave their assessment as originally provided (Burgman et al. 2011). Either decision requires justification. The sixth and final step requires assessors to provide possible reasons for why in their view initial assessments resulted in differences.

Information generated from this process was used to understand and classify the types of uncertainty, by treating the differences in assessment outcomes across assessors as types of uncertainty, and classifying them using the uncertainty typology of Regan et al. (2002). Sources of uncertainty were broadly classified as either epistemic or linguistic in origin; however, both broad categories of uncertainty were in some cases simultaneously possible as the cause of differences between assessment outcomes. Epistemic uncertainty is that associated with the knowledge of a system’s state and consists of six main types (measurement error, systematic error, natural variation, inherent randomness, model uncertainty, subjective judgment) (Regan et al. 2002). There are five main types of linguistic uncertainty associated with the omnipresent variation in our use of language (vagueness, context dependence, ambiguity, theoretical indeterminacy, under-specificity) (Regan et al. 2002). Other studies of uncertainty using this typology have modified it by, for example, including a third broad type of uncertainty (human decision uncertainty) which contain causes of uncertainty such as subjective judgment (Kujala et al. 2013). We chose to follow McGeoch et al. (2012) who used the original typology with minor adjustments for application to listing of invasive alien species, distinguishing between subjective judgment per se and subjective judgment due to a lack of knowledge, and between systematic error per se and systematic error due to a lack of knowledge. The rationale for this is akin to the dual pathway of forming a subjective probability (Burgman 2005), where people’s subjectivity stems from either a lack of, or despite, available knowledge.

Reasons for differences between assessments, i.e. the sources of uncertainty, were determined *a priori* by aligning these with the sources of discrepancies found in the alien listing process (McGeoch et al. 2012). Whilst some of the listing ‘errors’ were the same as those in McGeoch et al. (2012), others either needed to be re-interpreted for the assessment of impacts specifically or were not relevant for uncertainty associated with assessing impact. Also, using the initial results and the assessor’s judgments on why discrepancies occurred, additional reasons for disagreement specific to assessing impacts were identified. In total, we identified 12 sources of uncertainty in the impact assessment process and attributed each source to an uncertainty type (Table 1). Each species for which there was a difference between either mechanism or severity of impact, we then identified the reason(s) and their associated uncertainty. We did not assess the uncertainty associated with differences in the chosen confidence level due to its strong dependence on the chosen mechanism and severity.

**TABLE 1.**
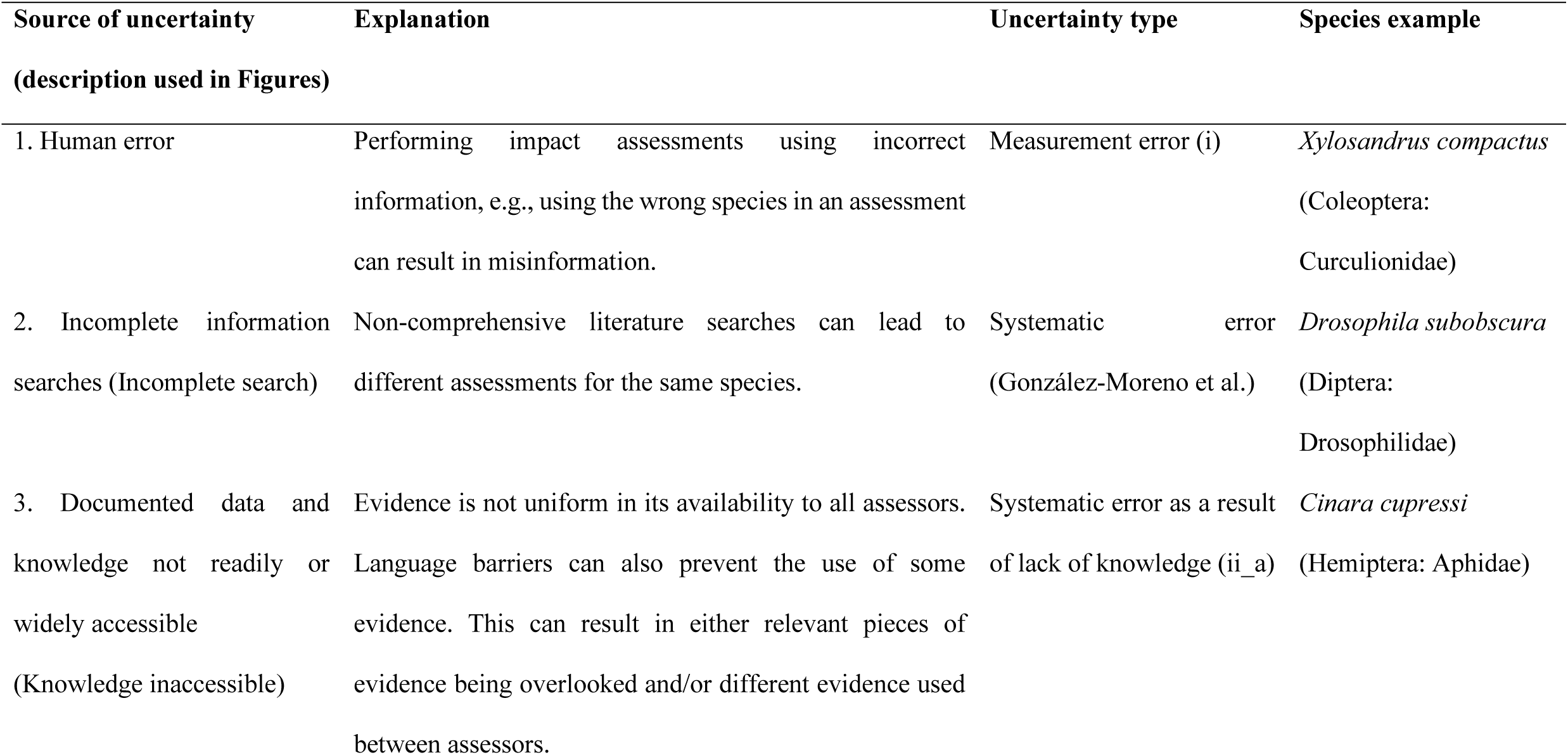
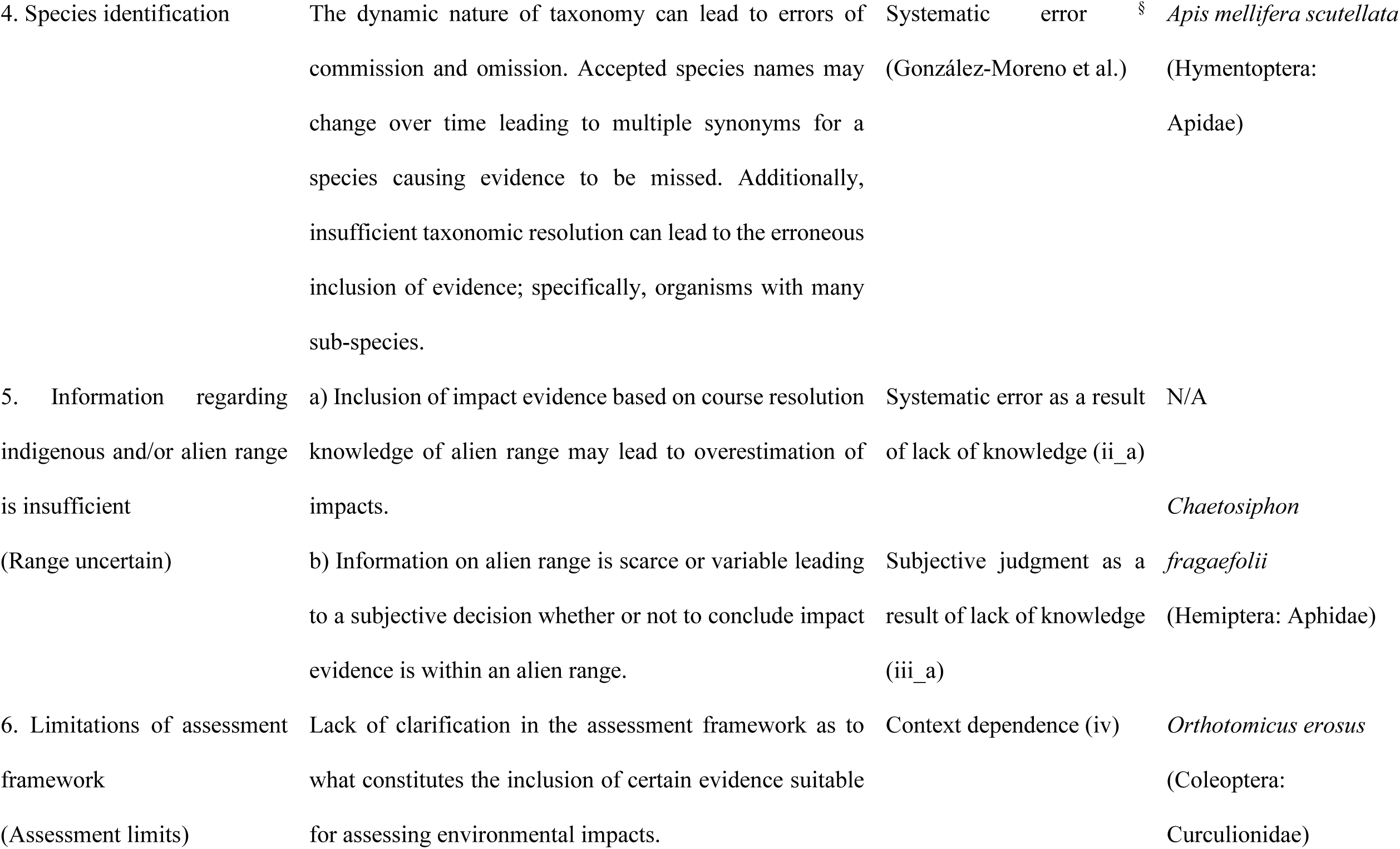
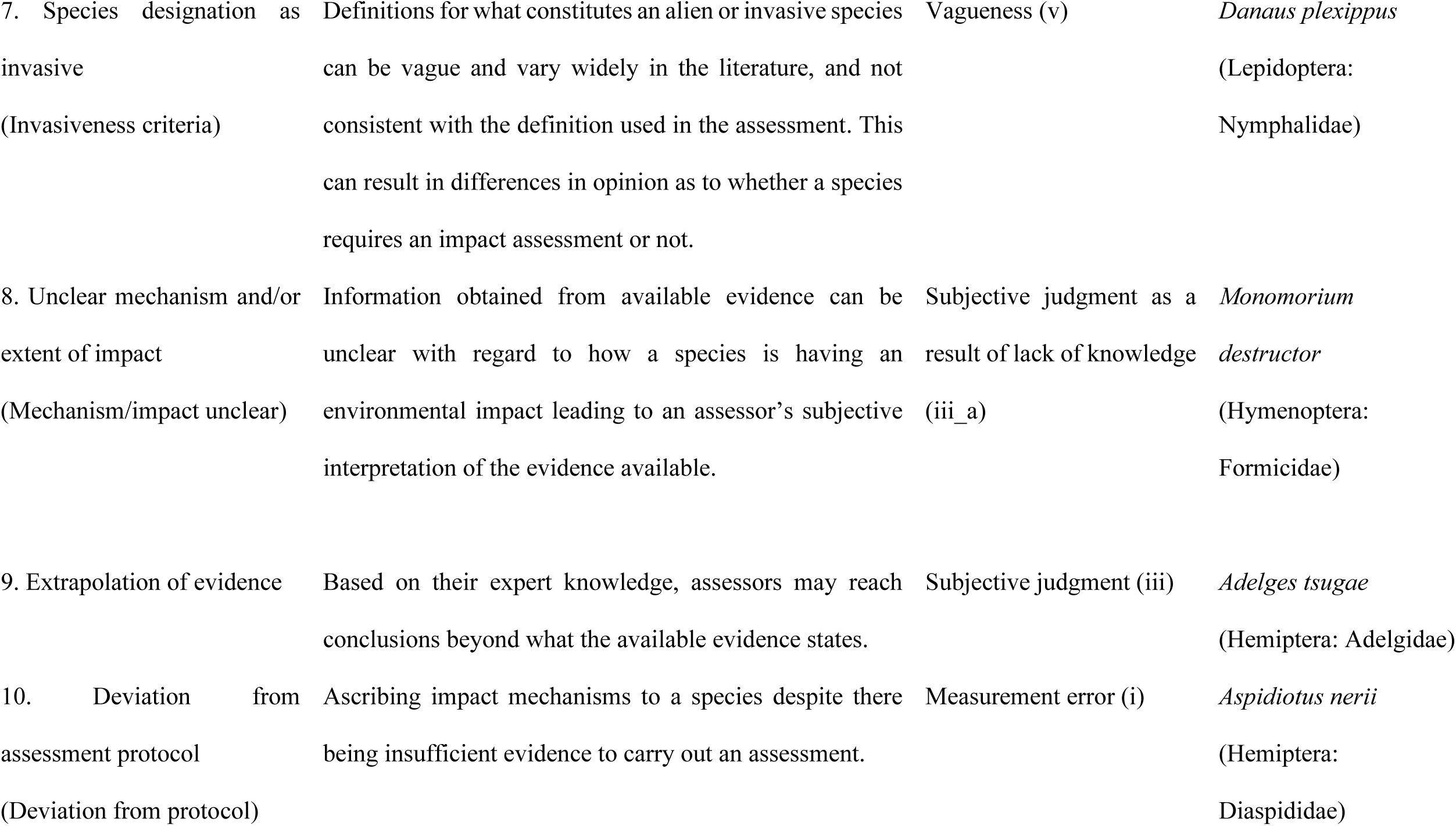
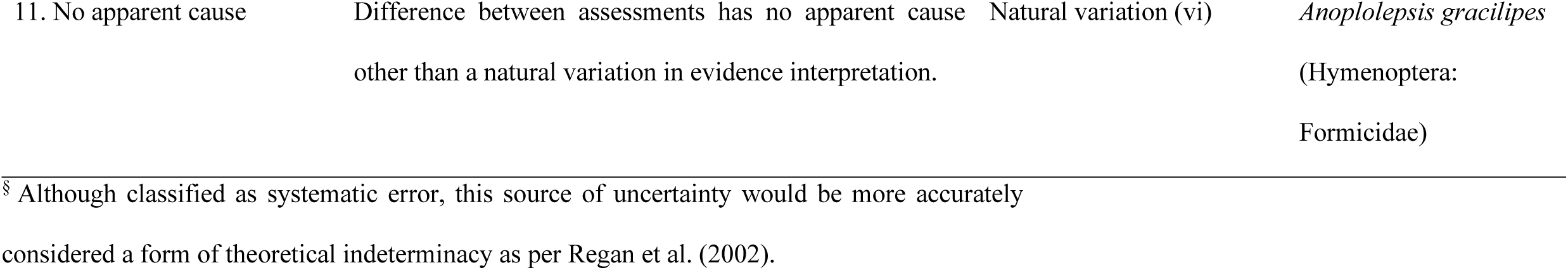
Sources, and types, of uncertainty that may lead to differences between independent assessors when performing environmental impact assessments. Species examples encountered during the process are discussed further in the text.

## RESULTS

First-round assessments led to agreement levels of between 32 and 44% for each of the three EICAT assessment components, i.e. mechanism, severity and confidence (Fig. 1). The mechanism of impact had the highest level of agreement, with 44% of species attributed with the same primary mechanism of impact by both independent assessors. The level of agreement however varied across mechanisms, as did the frequency with which each mechanism was attributed (Table 2; Fig. S2). Herbivory, competition and predation were most frequently identified (Table 2), and agreement on herbivory as the primary mechanism of impact occurred in 38% (*n* = 14) of instances. The remaining 8 mechanisms were less frequently attributed and were for the most part each either in agreement or disagreement (Table 2). For example, parasitism was attributed to 6 species, 5 of which were cases of agreement. Facilitation of native species was also attributed to 6 species, however, all 6 were cases that differed between assessors. Multiple (*n* = 14) instances occurred where the assessors agreed on a mechanism for a species, but one assessor also assigned additional mechanisms. The assignment of multiple mechanisms in EICAT occurs when the same level of impact severity is assigned to more than a single mechanism by an assessor for a given species. Taking a conservative approach here, these cases were considered differences in assessment outcome. After the second round of assessments, agreement on the mechanism of impact increased from 44 to 65% (Fig. S2). Little change was found among the individual mechanisms with a few exceptions; most notably herbivory (37 to 53%), other (6 to 40%), and transmission of disease (33 to 100%) (Table 2).

**TABLE 2.**
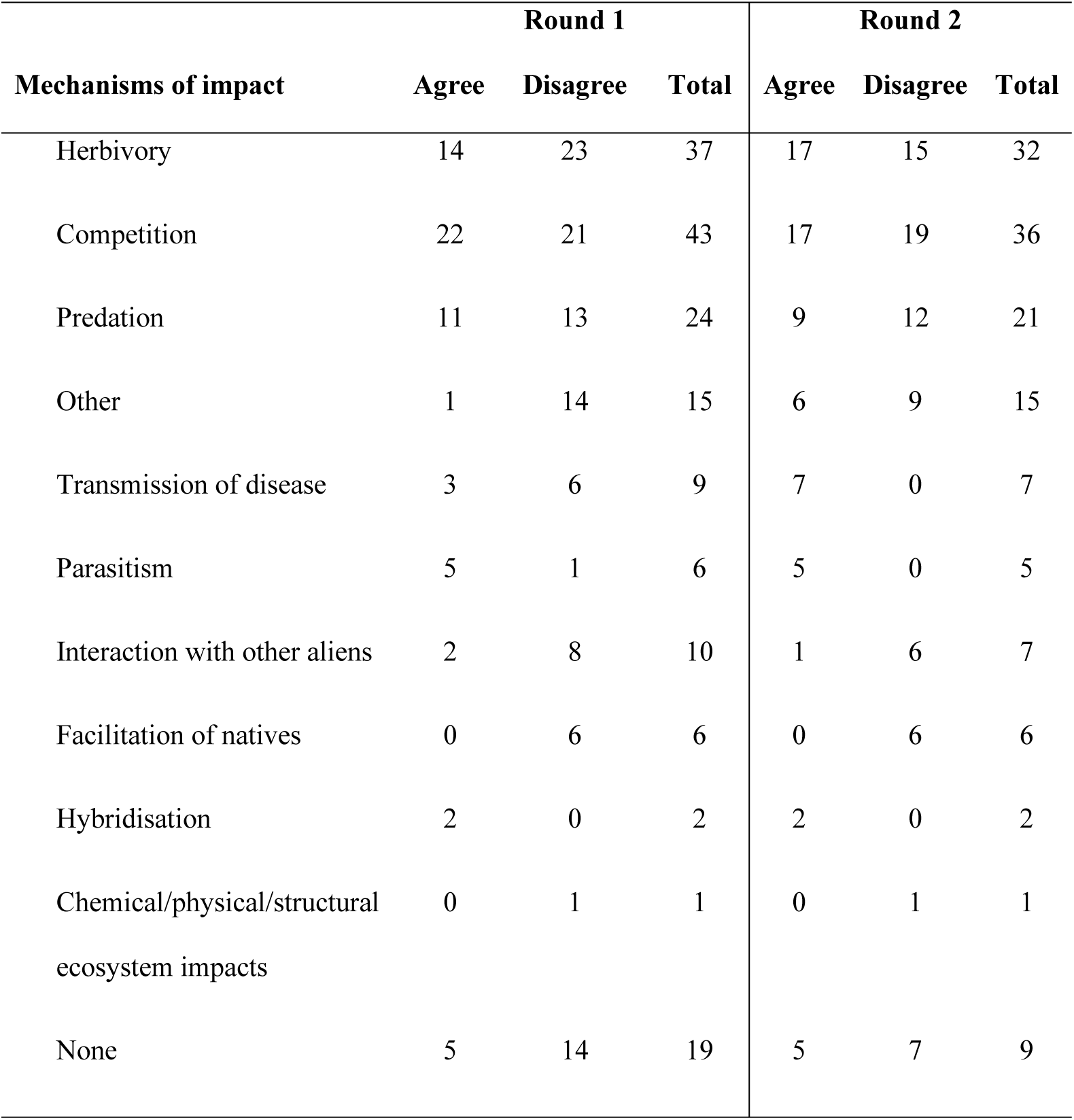
Instances of agreement and disagreement between assessors for mechanism of impact for both assessment rounds. (Note: As more than one mechanism can be attributed to a species, the instance totals within and between assessment rounds do not sum to 100.)

| Mechanisms of impact | Round 1 |  |  | Round 2 |  |  |
| --- | --- | --- | --- | --- | --- | --- |
|  | Agree | Disagree | Total | Agree | Disagree | Total |
| Herbivory | 14 | 23 | 37 | 17 | 15 | 32 |
| Competition | 22 | 21 | 43 | 17 | 19 | 36 |
| Predation | 11 | 13 | 24 | 9 | 12 | 21 |
| Other | 1 | 14 | 15 | 6 | 9 | 15 |
| Transmission of disease | 3 | 6 | 9 | 7 | 0 | 7 |
| Parasitism | 5 | 1 | 6 | 5 | 0 | 5 |
| Interaction with other aliens | 2 | 8 | 10 | 1 | 6 | 7 |
| Facilitation of natives | 0 | 6 | 6 | 0 | 6 | 6 |
| Hybridisation | 2 | 0 | 2 | 2 | 0 | 2 |
| Chemical/physical/structural ecosystem impacts | 0 | 1 | 1 | 0 | 1 | 1 |
| None | 5 | 14 | 19 | 5 | 7 | 9 |

**FIG. 1.**
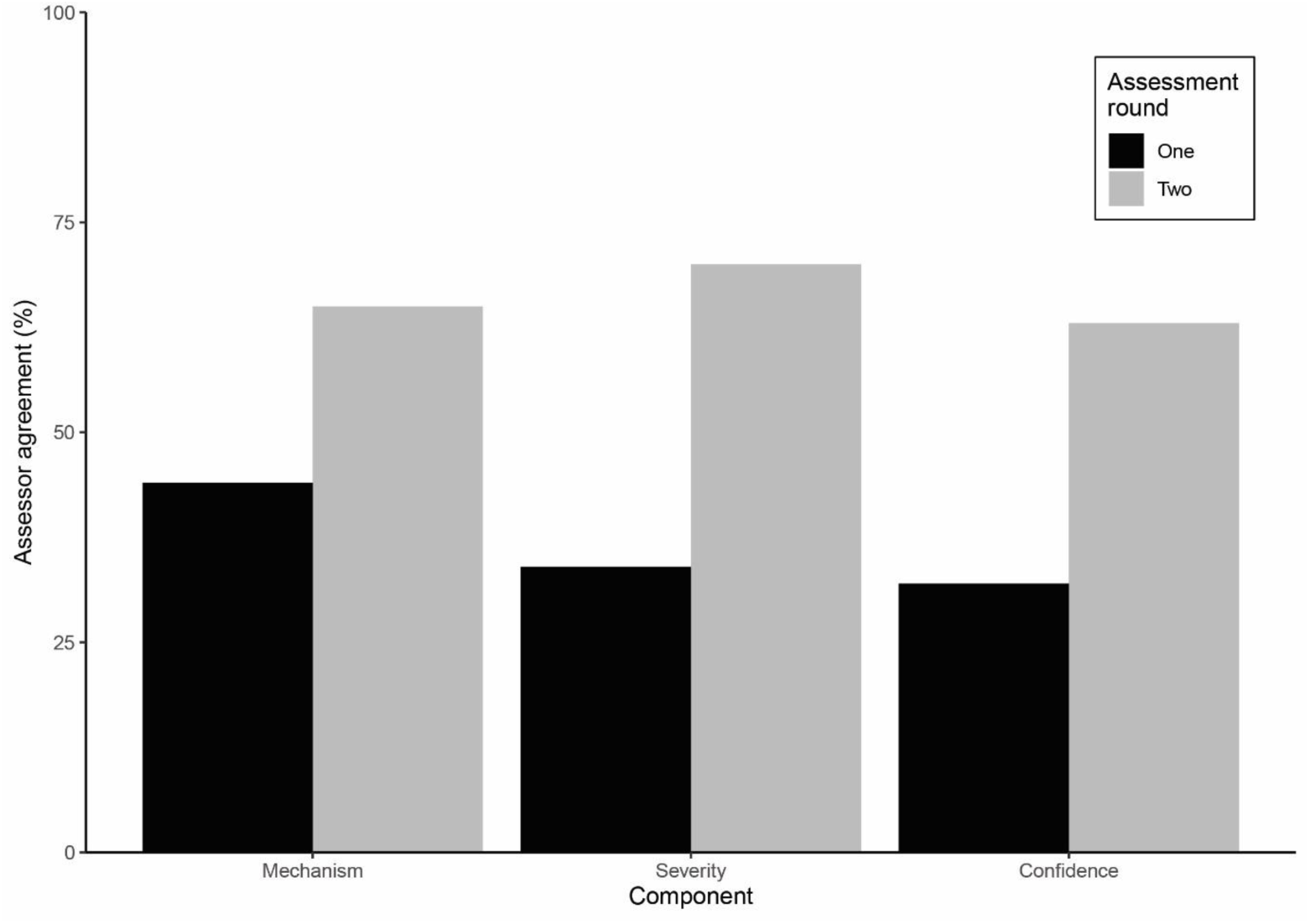
Agreement in environmental impact assessment outcomes between two independent assessors across two rounds of assessment. Agreement increased across all assessment components in the second round. The assessment of ‘severity of impact’ improved most and ‘mechanism of impact’ least across the two assessment rounds.

Agreement on impact severity in round 1 occurred for 34% of species, i.e. a species was assigned the same severity of impact by both assessors (assessment outcomes were as likely to agree as to differ, *χ*^*2*^ = 1.37, *df* =1, *P* = 0.24). Categories of impact severity varied in their levels of agreement (Fig. 2A). Data deficient (DD) was the impact severity classification with highest agreement (see Appendix S1: Table S1 for full details of impact severity assessments). Assessor agreement on impact severity increased from 34 to 70% across rounds. As in the first round of assessments, variation in agreement among severities of impact remained (Appendix S1: Table S1). Similarly, the distribution of agreement across the severity of impact categories differed little between assessment rounds (Appendix S1: Table S1). Assessors were more likely to agree than disagree (*χ*^*2*^ = 83.64, *df* = 1, *P* < 0.001) and most differences (22/30) between impact severity categories in the second round involved categories adjacent on the severity scale (Fig. 2B, Appendix S1: Fig. S1). For example, if there was difference in a moderate impact, it involved one assessor assigning either a minor (one category lower) or major (one category higher) impact (Fig. 2B, Fig. 3). Exceptions to this occurred for two species (*Diaphora citri, Tetropium fuscum*) in which one assessor assessed severity as minor and the other assessed it as major, and for six species (*Orthotomicus erosus, Solenopsis richteri, Chaetosiphon fragaefolii, Polistes chinensis, Rhyzopertha dominica, Thaumetopoea processionea*) in which one assessor assigned an impact severity and the other thought there was insufficient data to do so, or that there was no evidence of alien populations in the case of *T. processionea* (Fig. 3).

**FIG. 2.**
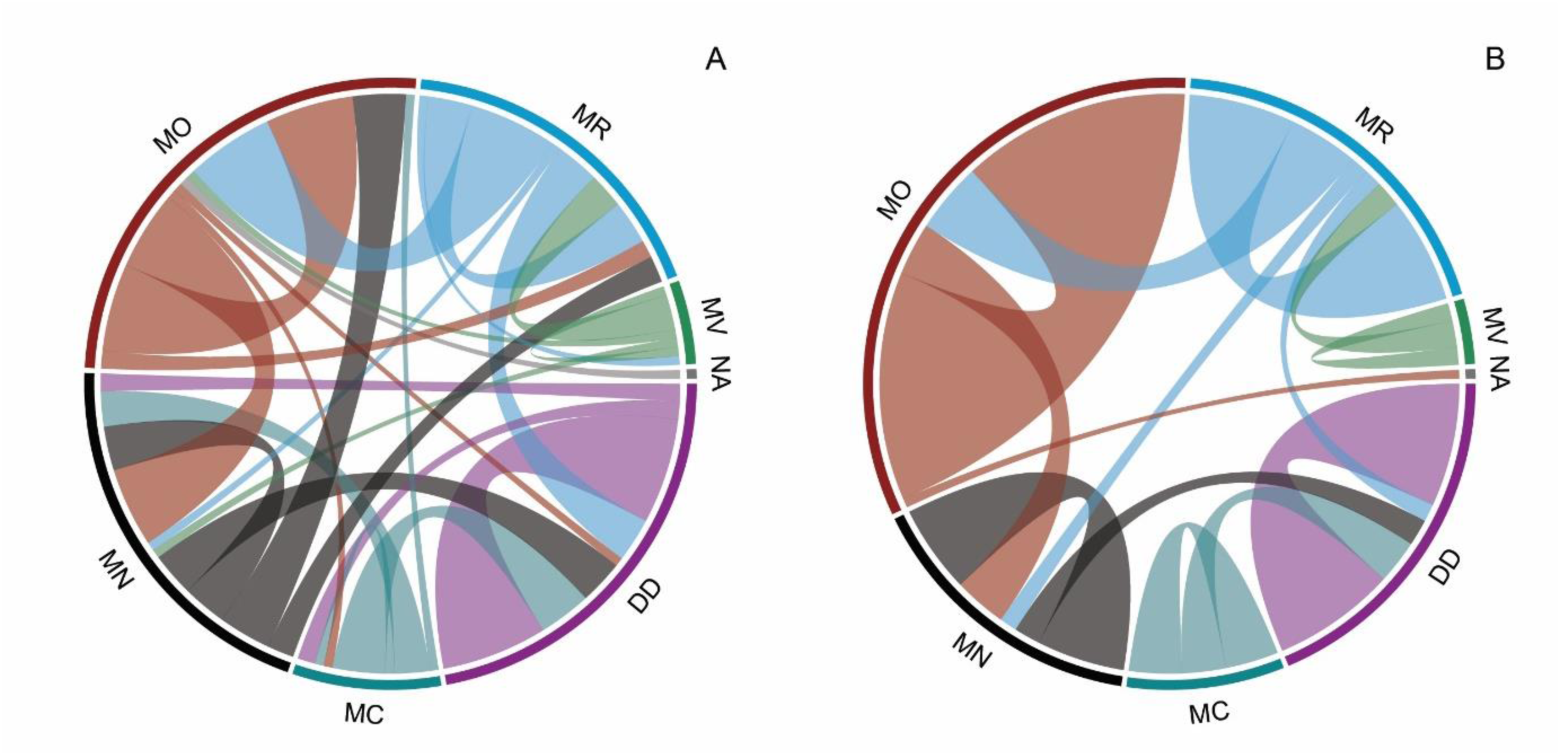
Chord diagrams showing the variation in agreement or disagreement in the assessment of severity of impact between independent assessors in each assessment round (A, B). The segment span width of a given impact severity category (MV (massive), MR (major), MO (moderate), MN (minor), MC (minimal concern), DD (data deficient) and NA (not alien)) is proportional to the number of species for which that severity was assigned by at least one assessor. The width of the links connecting segments reflects the number of species for which there was difference between assessors involving those specific impact severities. The low density of links in B compared to A reflects the increase in agreement across assessments, revealing also moderate followed by major and data deficient as the most frequently assigned categories.

**FIG. 3.**
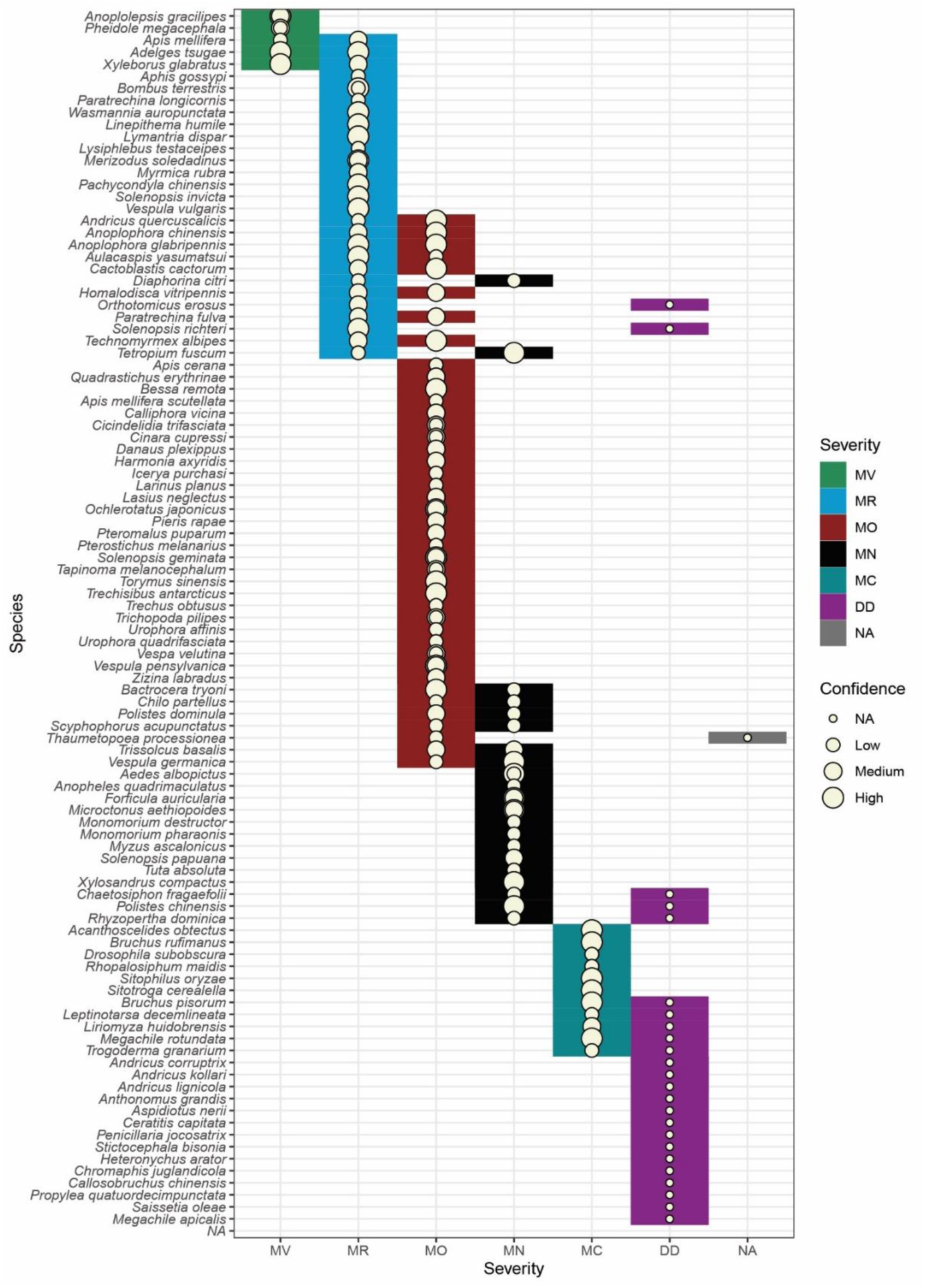
Final assessment results of impact severity and the associated confidence rating for all 100 insect species. Species for which there remained difference after both rounds of assessment are shown with two different coloured tiles in the row associated with them. Most instances of difference occurred at the interface between major and moderate, and between minor or minimal concern and data deficient. Category abbreviations: massive (MV), major (MR), moderate (MO), minor (MN), minimal concern (MC), data deficient (DD), not alien (NA).

All 11 possible sources of uncertainty identified *a priori* for assessor disagreement when performing environmental impact assessments (Table 1) occurred across the 100 species assessed. Most frequently, different use of reference material was part of, if not the entire cause of difference in a species assessment, closely followed by limitations of the assessment framework and the mechanism or extent of impact being unclear (Fig. 4). However, the frequency of these sources of uncertainty differed depending on the assessment component (mechanism or severity). For example, although extrapolation of evidence and deviation from assessment protocol occurred similar numbers of times (14 and 12 times, respectively), all but one of the former cases were related to differences in severity, whilst all instances of the latter were related to differences in mechanisms (Fig. 4). Systematic error was the uncertainty type that occurred most frequently, followed by subjective judgment as a result of lack of knowledge and context dependence (Fig. 5). Again, assessment components differed in the frequency of uncertainty types, most notably the difference in frequency of subjective judgment. Of the 14 instances where this was the type of uncertainty was involved, 13 were related to severity of impact.

**FIG. 4.**
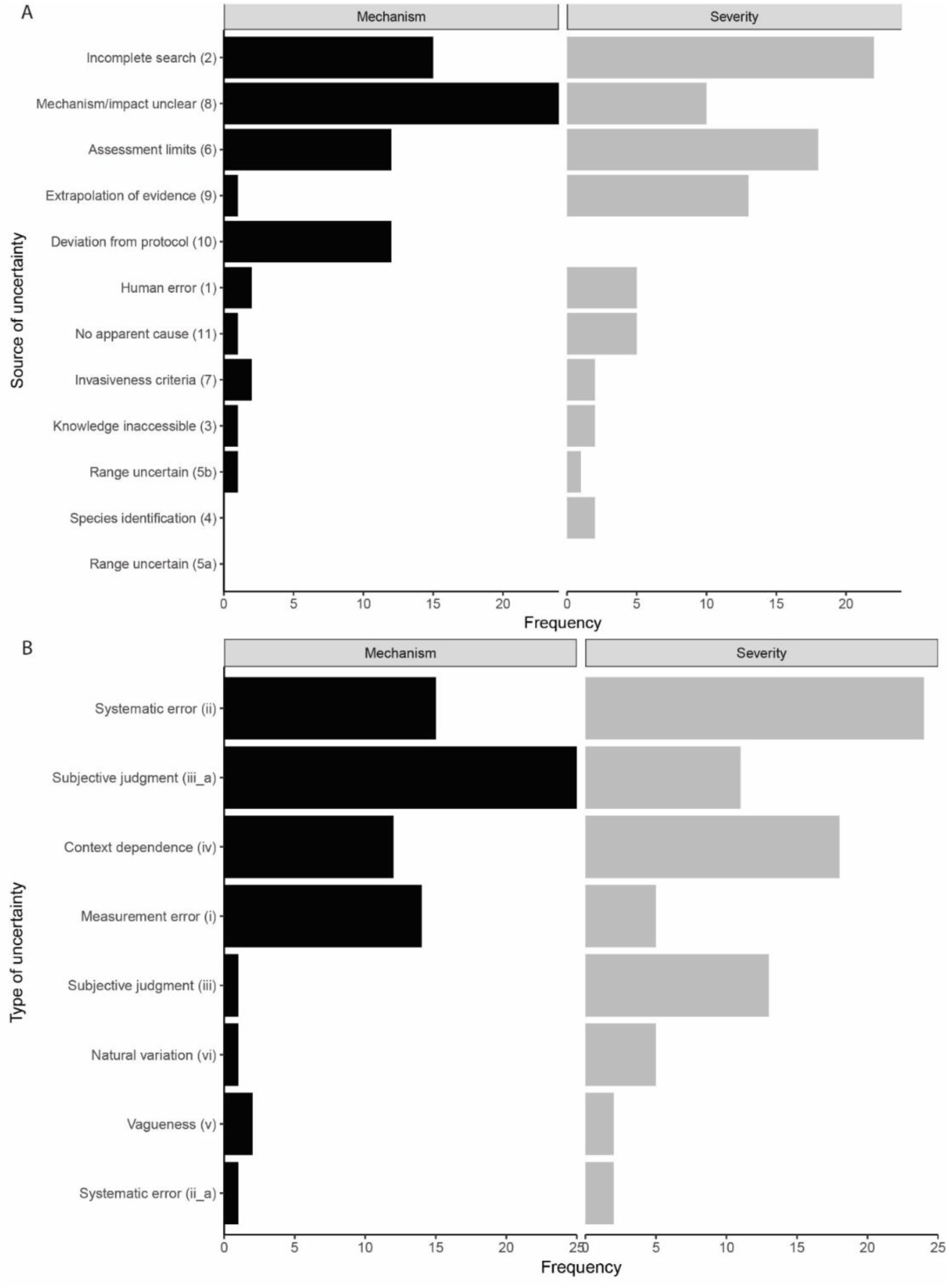
Frequency with which (A) sources of uncertainty and (B) types of uncertainty varied within and between impact assessment components (mechanism and severity of impact). Bar lengths highlight the dominant sources (A) or types (B) of uncertainty. The number and Roman numerals alongside the sources and types of uncertainty are those used in Table 1, and ii_a and iii_a are the sub types related to lack of knowledge.

**FIG. 5.**
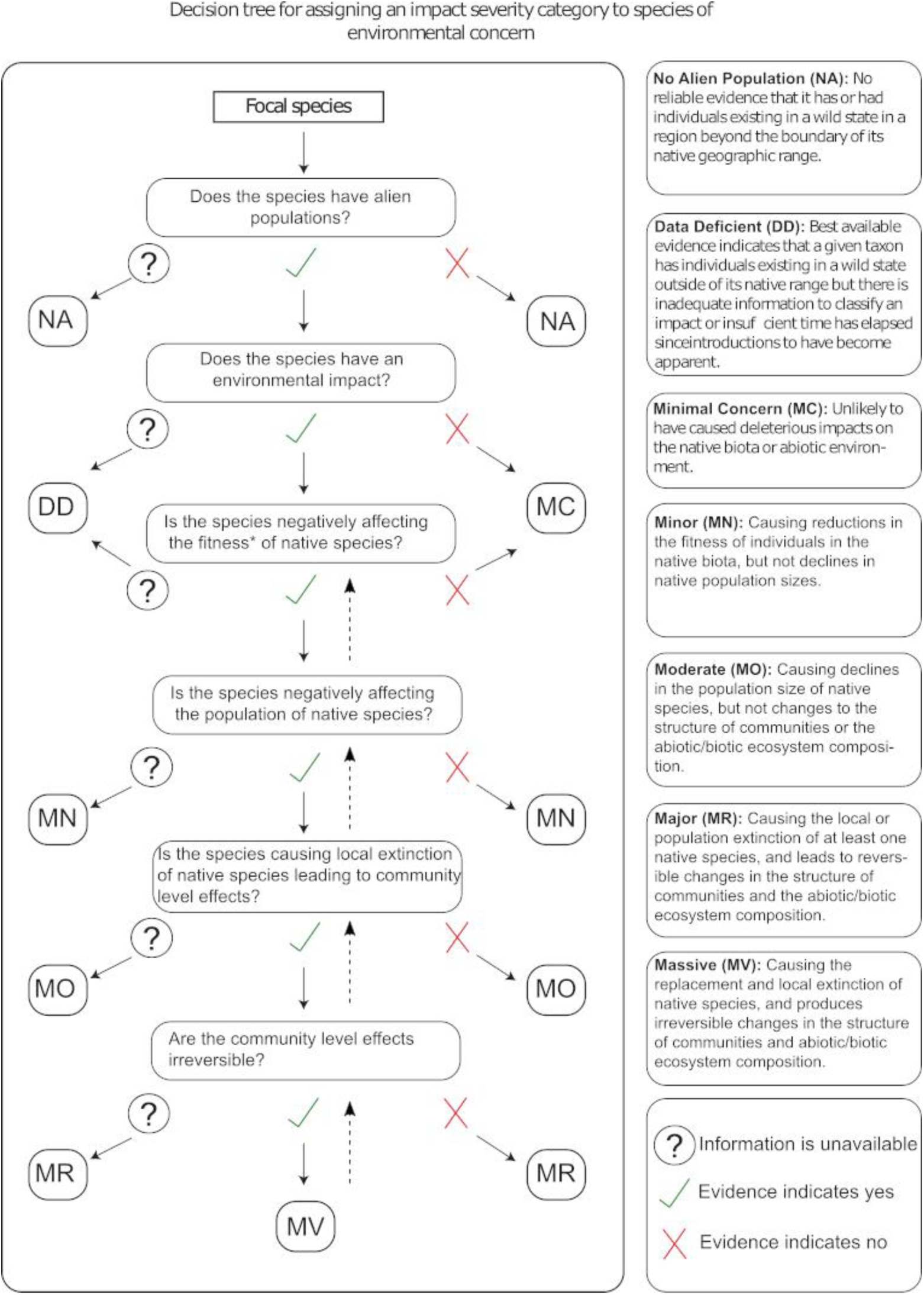
Decision tree for improving the transparency and repeatability of assigning EICAT impact severity categories. This framework is based on the assumption that the different categories (MC to MV) are hierarchically related, i.e. a negative effect at one level necessitates a negative effect at the level below it (dashed arrows) and may mean existence of an impact at a higher impact severity level, although no evidence may yet be available to support designation of this higher level of impact. Under this assumption there can often be two pathways by which an impact category can be assigned, with the followed path dependent on information availability. For example, in the case of the two pathways for assigning a moderate (MO) impact; if there is information to support the conclusion that there is a negative effect on the population size of one or more native species, but no effect on community structure, then the appropriate category to assign is MO. Alternatively, there may be information on native population level negative effects but no information (lack of evidence) available on the effects at the community level, in which case the category assigned would also be MO. In general terms, these two alternative pathways represent ‘evidence of no impact’ (right) and ‘no evidence of impact’ (left). *In the most recent version of the EICAT implementation guidelines (https://www.iucn.org/theme/species/our-work/invasive-species/eicat), the term fitness has been replaced by performance.

Final assessments revealed evidence of environmental impact was inadequate or unavailable for 14 species (Fig. 3), i.e. where both assessors agreed on the species being data deficient (Appendix S2: Table S1). Including the additional 10 species where assessors disagreed on data availability equates to approximately 25% of alien insects assessed as data deficient by at least one assessor. These species were distributed across five orders: Coleoptera (*n* = 9), Diptera (*n* = 2), Hemiptera (*n* = 5), Hymenoptera (*n* = 7) and Lepidoptera (*n* = 1). Of those species for which environmental impact information was available and assessors agreed on impact severity (*n* = 56), 25 were Hymenoptera. This order was most frequently represented in each impact severity category, except for minimal concern where there were no Hymenoptera. It was also the only order for which there was agreement on a massive impact, i.e. for both the yellow crazy ant (*Anoplolepsis gracilipes*) and the Big-headed ant (*Pheidole megacephala*). Few species were attributed this highest level of impact, and only five species were assigned this category by at least one assessor (Fig. 3). Considering only those species for which there was agreement on severity and excluding those that were data deficient, the number of species across impact severity categories approximates a symmetric distribution with moderate (*n* = 26) the most commonly attributed category (Appendix S1: Table S1). Including instances of difference alters the distribution of impact severities, which reflects the uneven frequency with which each category was associated with assessment differences.

Agreement on impact mechanisms was most common for herbivory (*n* = 17) and competition (*n* = 17) (Appendix S1: Table S1). Hemiptera were somewhat of an exception and here transmission of disease occurred as often as herbivory. Although these two mechanisms were most frequent, all orders except one (Dermaptera) were associated with at least four mechanisms of impact (Appendix S1: Fig. S2). This declined slightly when considering only instances of agreement, with Lepidoptera decreasing to two attributed mechanisms.

## DISCUSSION

Here we identified the types of uncertainty in impact assessment for alien species, and which of these types and their sources are amenable to reduction. Using insects as the model taxon for this exercise, we provide the impact severity results for 100 alien insect species of environmental concern with associated estimates of uncertainty. Uncertainty in the environmental impact assessment process for alien species can lead to independent assessments arriving at different conclusions for the same species, or single assessments resulting in biased outcomes. Neither of these alternatives is desirable when decision makers rely on such information to guide action and investment. Examining alternative outcomes for individual species provided insight on the type and extent of uncertainty involved, and potential solutions for reducing it. We found that following the assessment guidelines resulted in relatively high levels of disagreement between independently conducted assessments, with the reasons for these differences associated with 11 different sources and six different types of uncertainty. However, levels of agreement increased substantially following discussion between assessors for particular species and discussion of the reasons for the uncertainty involved. This process enabled us to identify several practicable recommendations for reducing the uncertainty associated with assessing the impact of alien species.

The process we used to select species to be assessed maximised the chance of information being available for each species, because well-known taxa were selected deliberately. Indeed, information was available for 75 (or 86% if one excludes differences where only one assessor assigned data deficient) of the insect species assessed. Around 20% data deficient species is low by comparison with the outcomes of EICAT assessments conducted to date on other taxa (71% of alien birds (*n* = 296/415) (Evans et al. 2016), 62% of amphibians (*n* = 65/105) (Kumschick et al. 2017b), and 85% of bamboo (*n* = 115/135) (Canavan et al. 2019), although 24% of gastropods were classed as DD (*n* = 8/34) (Kesner and Kumschick 2018)). This outcome (20%) is therefore likely a best-case scenario for the level of data deficient species in impact assessments for alien insects, and may be expected to increase as more alien insects are assessed.

Although straight-forward comparisons between insects and other taxa are complicated by differences in the number of species assessed, the range of high impact insect species is similar to other assessed taxa. The 100 insect species assessed included 14-29 % (by both or single assessors) species that were assigned a severity category of massive or major. This is compared with 15, 23, 8, and 10 % of gastropod, amphibian, bird and bamboo species respectively (Evans et al. 2016, Kumschick et al. 2017b, Kesner and Kumschick 2018, Canavan et al. 2019). Again, because we deliberately selected well-known invasive alien species, this percentage of invasive alien insects with severe environmental impacts can be expected to decline as more insect species are assessed.

### Types of uncertainty and their sources

Agreement on severity of impact increased from 34 to 70% following the second round of assessments, suggesting that this assessment component is quite amenable to a reduction in uncertainty. Nonetheless, this also means that assigning severity of impact encompasses more uncertainty than assigning a mechanism of impact. As with mechanisms, incomplete information searches (systematic error, Table 1, 2, ii) were a common cause of difference, and the most frequent reason for why independent assessors assigned different impact severities. There are several possible explanations for why different pieces of evidence were selected for use, not least of which are different interpretations of what constitutes evidence appropriate for inclusion in an environmental impact assessment (i.e. limitations of assessment framework, Table 1, 6, iv). Indeed, this was the second most common source of difference in impact severity assigned to species and may to some extent be particular to the assessment of insects.

One of the more common sources of difference in assigning a mechanism of impact was a lack of clarity in the publications used as evidence for how an alien species was negatively affecting the native environment. In such cases, assessors were required to make a subjective interpretation about what mechanism was most likely, based on the context of the study. Similarly, it was often difficult to differentiate between two mechanisms, for example competition and predation, where the evidence suggested that either choice was valid. This led to situations where either one assessor chose competition and the other predation, or where one assessor provided both competition and predation because on the basis of the evidence they were not able to distinguish between them. For example, the evidence of environmental impact for the harlequin ladybeetle (*Harmonia axyridis*) (Roy et al. 2012, Masetti et al. 2018) and the Argentine ant (*Linepithema humile*) (Menke et al. 2018) often inferred the impact mechanism could be competition or predation. In situations such as this where the evidence itself is unclear on the mechanism of impact, it is difficult to avoid differing subjective interpretations by independent assessors (i.e. subjective judgement as a result of lack of knowledge, Table 1, 8, iii_a).

Another common reason for differences between assessors on impact mechanisms, was the use of different lines of evidence (systematic error, Table 1, 2, ii). Multiple reasons for this are possible. First, for many insects the quantity of literature is large, and a literature search can return thousands of studies for a single species (for example, some species assessed had search returns in the order of 14 000 publications). A large literature on a species increases significantly the time needed to conduct an assessment. Simple oversight can result in mistakenly missing, misinterpreting or including a certain piece of evidence (increasing the chance of human error and incomplete information searches as the source of uncertainty (Table 1, 1i, 2ii)). For example, differences in the first round of assessments for the shot-hole borer (*Xylosandrus compactus*) occurred because of the accidental use of evidence pertaining to a congeneric species (Table 1). By contrast, the Mediterranean pine beetle (*Orthotomicus erosus*) assessments differed as a result of different interpretations of the EICAT guidelines (Table 1, 6). Both examples become sources of uncertainty for reasons other than simply incomplete information searches. This shows how uncertainty can compound during the assessment process, starting with one uncertainty type (in this case measurement error) that leads to another (context dependence). However, this also implies that by addressing some of the more manageable sources of assessor difference, uncertainty propagation could feasibly be reduced (see recommendations below).

Even with definitions for each severity category (Hawkins et al. 2015), assigning impact severity appears to be a decision open to greater interpretation than assigning impact mechanisms. One explanation may be the naturally unbounded nature of the relationship between successively more severe impact categories, where even with definitions for each impact severity category there is inevitably scope for subjective interpretations of degree. For example, an alien species with a minor impact is one that negatively affects the fitness (or performance, see Fig. 5) of a native species. However, a decline in fitness of individuals is likely to lead to a decline in population size, which constitutes the next most severe impact category (moderate). The difference between a negative impact on the fitness of individuals in a population and a population-level effect is a matter of degree (Ricciardi et al. 2013, Blackburn et al. 2014). Thus, if we assume a given impact severity begets the next most severe category, then there are two alternative pathways to arrive at a given severity category, i.e. either via *evidence of no impact* or via *no evidence of an impact* at the next most severe category (use of the decision tree in Fig. 5 enables these alternatives to be distinguished). For example, an assessor may conclude that a given species is having a MN impact either because (1) there *is evidence* it is *not causing* population declines, or (2) because there is *no evidence* it *is causing* population declines. For example, Hadley and Betts (Hadley and Betts) found that a proposed lack of negative consequences of habitat fragmentation on pollination dynamics was a result of the absence of evidence, and not evidence of an absence of such consequences. Hawkins et al. (2015) state that an impact severity category is assigned if there is no evidence for the next most severe category, implying the absence of evidence case (2 above). However, explicitly distinguishing between these two scenarios is valuable, even if the former occurs rarely by comparison, because it informs the strength of evidence available and confidence placed in it. Specifying such reasoning during assessments by making the decision process explicit (as shown in Fig. 5) could reduce uncertainty, as well as make the process more readily repeatable by specifying the basis for the final decision, i.e. that there was, or was not, evidence for the next most severe category (Fig. 5).

Assessing the environmental impact of alien insects may bring with it added complexity not experienced with other taxonomic groups for which environmental impact assessments have been published to date. Primarily, this complexity, and a large source of uncertainty, concerns the many environmental settings in which invasive insects can have a negative impact (Kenis et al. 2009, McGeoch et al. 2015). This can add uncertainty to decisions on which evidence to include (context dependence, Table 1, 6, iv). For example, the Mediterranean pine beetle has been unintentionally introduced to multiple countries, most likely via the timber trade (Brockerhoff et al. 2006, Haack 2006). It is known to cause damage to conifers, particularly pines, often leading to tree mortality (Sangüesa-Barreda et al. 2015). However, all such evidence of a negative impact inside its alien range takes place within managed forestry plantations (Stephens and Wagner 2007)). As such, both assessors for *O. erosus*, who differed in opinion as to which evidence to include, came to entirely different conclusions even after the second round of assessments (data deficient versus massive) (Haack 2004). Similar discrepancies occurred in the assessments of the Citrus long-horned beetle (*Anoplophora chinensis*) and the Asian long-horned beetle (*A. glabripennis*). Much of the damage caused by these species in their alien range occurs within urban environments, negatively affecting trees occupying green spaces (Sjöman et al. 2014). Given the urban setting, one assessor concluded the species to be data deficient whilst the other assigned it major impact severity. This assessment changed after the second round of assessments following the decision that, although the evidence is of impacts occurring in an urban environment, if the species being affected are native then such evidence is valid for inclusion and warrants an impact classification (Hawkins et al. 2015).

The large majority of impact research on alien insects is socio-economic, with much of the evidence for insects related to negative impacts to agricultural crops (Bebber et al. 2014), forestry plantations (Wingfield et al. 2015), building structural integrity (Mahapatro and Chatterjee 2018), and human health (Juliano and Philip Lounibos 2005). Some of these systems nonetheless represent grey areas (ecological systems requiring clarification) in terms of their relevance to environmental impact assessment. For example, forestry plantations established for the purpose of timber harvest could arguably be considered ‘natural’ (i.e. of environmental relevance in the context of EICAT) or agricultural, depending on the tree species planted and if it is native or not (Chazdon et al. 2016). Similar arguments can be made for agricultural crops and urban environments, depending on the extent to which native biodiversity or native relatives within them are impacted by the alien insect that thus poses a potential risk to native biodiversity more widely. Without formal direction and clarification at the outset of the assessment process, differences based on subjective interpretations of ecological systems suitable for inclusion are inevitable, albeit avoidable. Inclusion of an alien insect species in environmental risk assessment should, we argue, be based on *what* is being negatively affected more so than *where*, i.e. the context within which the impact is occurring is less relevant in some cases.

A different situation arises for insect species that may arguably never negatively affect the native environment, and this assumption led to some species being assigned minimal concern despite the absence of evidence (subjective judgement, Table 1, 9iii). Two groups of alien insects, given their life histories, may lead assessors to assign Minimal Concern without explicit evidence (i.e. rather than data deficient). The first group are those species that are primarily stored product pests. For example, the bean weevil (*Acanthoscelides obtectus*) is an economically important pest of legumes (Soares et al. 2015). Whilst damaging, its impact almost entirely occurs on stored products, and it is in essence a species solely of economic concern with a very low likelihood of having an environmental impact. The second group are monophagous insects that only attack and exist in agricultural crop environments and/or have a long history of no evidence of environmental impact. The broad bean beetle (*Bruchus rufimanus*) is a widespread crop pest, specifically targeting the faba bean (*Vicia faba* L.) (Clement et al. 2002). Given the long history of evidence for species such as this only being of concern for specific crop types, one could conclude that it is of minimal environmental concern. However, based on the lack of explicit evidence of no effect these cases would otherwise be classified as data deficient. Such conclusions are an instance of using absence of evidence as evidence of absence; generally undesirable but logically valid under certain circumstances (Sober 2009). The decision to classify a species as minimal concern for environmental impact without evidence, however, should occur outside of the impact assessment process in the form of additional information that may complement the final assessment outcome, as suggested for accommodating expert input (see recommendations below).

The complexity of assessing environmental impacts of alien insects extends beyond the broad range of ecological systems that they invade to the range of mechanisms of impact they can have (McGeoch et al. 2015). Across the alien insects assessed, at least one instance of each of the 11 possible mechanisms of impact was recorded, albeit with disproportionate frequencies. However, one of the most frequent mechanism categories was “other”, i.e. the evidence suggested a mechanism not currently listed under the EICAT protocol. For example, the gall wasp (*Andricus quercuscalicis*) was assessed as having an impact by altering the sex ratio of native species (Schönrogge et al. 2000); a mechanism that does not easily fit within any of the other mechanism categories. The Asian honey bee (*Apis cerana*) robs food stores of native species (Hyatt 2012) and was noted as possibly fitting under multiple mechanism categories. Differences in mechanism choice between assessors involving the use of ‘other’ often also involved the mechanism ‘interaction with other alien species’ or ‘facilitation of native species’. This may imply that “other” is often used to denote some form of indirect interaction that is associated with a negative environmental impact. Insects are frequently involved in such indirect interactions (Kaplan and Denno 2007, Dáttilo et al. 2016) and alien insect species are no different (McGeoch et al. 2015). If the outcome of an assessment is the assignment of “other” for the mechanism of impact, then the assessor should provide a concise description of what they interpret the mechanism to be. Such information would be useful not only for the assessment itself, but also for potentially updating the framework to include a new mechanism of impact (IUCN 2019). This could subsequently reduce uncertainty associated with impact mechanism decisions.

Not all sources of uncertainty have equally significant consequences for the outcome of an assessment. For example, deviation from assessment protocol (measurement error, Table 1, 10, i) in some instances was simply the situation where one of the assessors, although assessing a species as data deficient did actually provide a mechanism of impact, rather than assigning “none” (e.g. Oleander scale (*Aspidiotus nerii*) Table 1). This tended to occur when the assessor was familiar with the species and providing an educated judgement as to the most likely mechanism of impact. For example, Lepidoptera are likely to affect via herbivory and carabids by predation (or competition). Discrepancies of this kind are unlikely to have any serious consequences for species prioritization, and by extension resource allocation, and can be readily avoided. Extrapolating evidence to assign an impact severity beyond that provided by the available evidence is, however, more problematic (i.e. subjective judgement, Table 1, 9, iii). This was one of the more frequent causes of uncertainty associated with impact severity. Subjective judgment of this form stemmed primarily from the expert knowledge of those performing assessments on species with which they are particularly familiar. For example, impact severity for the Hemlock woolly adelgid (*Adelges tsugae*) differed as a result of this source of uncertainty (Table 1). Extended familiarity with a given species may reduce objectivity leading to a more liberal, but not necessarily wrong, interpretation of impact severity. This subjectivity can result in an assessment at odds with an assessor who is less familiar with the same species, but who therefore bases their assessment on a more objective interpretation of the evidence. We suggest that this source of uncertainty could be reduced by explicitly incorporating, within the assessment framework, opportunity for expert opinion to be provided in addition to and alongside the assessment based on published evidence alone (Morgan 2014); see Recommendations below).

Most types of uncertainty encountered when assessing the environmental impacts of alien species were epistemic in nature, a result similar to other studies on the presence and effect of assessment uncertainty (McGeoch et al. 2012, Kujala et al. 2013). The disproportionate occurrence of epistemic sources of uncertainty, however, does not downplay the importance of linguistic uncertainty (Table 1 7, v). Historically overlooked, this pervasive type of uncertainty can be influential, problematic, and difficult to reduce (Regan et al. 2002, Burgman 2005, Carey and Burgman 2008). This is particularly true in invasion biology where variation in definitions and their use have hindered progress in the field (Verbrugge et al. 2016, Courchamp et al. 2017). Crypticity, an aspect of biological invasions that can lead to underestimation of impacts, is also in part affected by linguistic uncertainty due to the dynamic nature of taxonomy (Jarić et al. 2019). Regan et al. (2002) characterised this as a form of theoretical indeterminacy where a term, or in this case a species name, will not necessarily retain its meaning into the future. Advances in molecular biology may unveil previously unknown species complexes or reveal that specific sub-species that are in fact alien to a given region (Jarić et al. 2019).

### Recommendations for reducing uncertainty in impact assessments for IAS

Certain sources of uncertainty are unavoidable and their presence and potential impact on assessment outcomes can only be acknowledged (Humair et al. 2014), whereas other sources can potentially be reduced. Based on a structured process for identifying sources of uncertainty, we recommend the following six steps to reduce uncertainty in impact assessments for alien species of environmental concern. While we used insects as a model taxon, the recommendations below are taxon agnostic.

1. The dissemination of information derived from risk analysis, and indeed impact assessments, is often overlooked (Vanderhoeven et al. 2017). Not only is it important to communicate the conclusions of impact assessments, such as the EICAT classification, but also to ensure the degree of uncertainty is transparently and effectively shared with end-users. There is a need to develop and implement risk communication and to ensure tools adopted are appropriate to the target audience. The IUCN Red List is used as a communication tool in many different contexts and EICAT could benefit from exploring the most effective approaches employed. Particular attention should be given to transparent communication of uncertainty to increase understanding of risk by relevant stakeholders.
2. For an impact assessment to convey the most accurate representation of the evidence, assessments should be performed as objectively as possible. Although perhaps self-evident, it is reasonable to assume that some assessors may use their expert knowledge of a certain species to extrapolate beyond available evidence. To accommodate this valuable information, assessments should include within them the opportunity for assessors to provide their expert judgement, in addition to but separate from the objective results of the assessment itself. This could allow what is analogous to uncertainty bounds around the final assessment outcome (e.g. as portrayed in Fig. 3).
3. Many assessment differences involving the use of “other” also involved mechanisms indicative of indirect or higher order interactions. As we found here, indirect interactions can involve facilitation of native species, as well as facilitation *by* native species in the establishment of introduced organisms (Northfield et al. 2018). Use of any attribution of “other” as mechanism of impact, as was frequent in this study, should always be accompanied by a description of the impact mechanism. This information would not only enrich the impact assessment but would also assist in future developments of the assessment framework to include new mechanisms of impact should similar descriptions be repeated over time.
4. An unambiguous definition for what qualifies as data deficient for environmental impact should be established to reduce misinterpretation about what evidence to include in an assessment. Insects have a negative impact in several different systems, including agriculture, forestry, and urban environments. Emphasis should be placed on *what* species is being negatively affected in addition to *where* the impact is taking place. In other words, the environmental context of the evidence should be specified and used as part of the rationale for including or excluding a given piece of evidence.
5. Use of a detailed decision tree to harmonise and capture the decision making process (such as Fig. 5), would improve the transparency and repeatability of assessments, and formally distinguish between ‘evidence of no impact’ and ‘no evidence of impact’, and therefore improve the rigour of assigning confidence to each decision.
6. Systematic review protocols should be explicitly integrated into the impact assessment guidelines and literature searches performed using established protocols such as the Guidelines and Standards for Evidence Synthesis in Environmental Management (Pullin et al. 2018, Siddaway et al. 2019). One of the more frequent sources of uncertainty was the use of different literature by assessors. Differences in evidence accrual procedures has also been identified to affect comparability of various impact assessment frameworks (Strubbe et al. 2019). Following such protocols would allow transparency and repeatability of the literature search process and should reduce the occurrence of different literature use between assessors for the same species. In addition, given the pervasive uncertainty associated with taxonomy (Pyšek et al. 2013), all known synonyms of the accepted scientific name of a species should be included in the search terms, thus ensuring the inclusion of historical information.

Incorporating the above recommendations is likely to increase the accuracy, transparency, and reliability of the alien insect impact assessment process and outcome. Regardless of the type (epistemic or linguistic), being aware of the potential sources of uncertainty in alien species impact assessments, and to a greater extent assessment processes in general, is important if we are to reduce uncertainty as much as possible and base our management and prioritisation decisions on the most accurate and reliable information available (Hamel and Bryant 2017, Latombe et al. 2019). This is particularly relevant given the current Intergovernmental Science-Policy Platform on Biodiversity and Ecosystem Services (IPBES) assessment on invasive alien species and their control (IPBES 2018).

## ACKNOWLEDGEMENTS

We acknowledge financial support from the Invasive Species Council, the Ian Potter Foundation, Monash University, the Australian Department of Agriculture, and the Queensland Department of Environment. We thank Andrew Cox, Carol Booth, and Rebecca O’Connor, for their comments. MAM acknowledges support from the Australian Research Council (DP200101680). DAC acknowledges support from an Australian Government Research Training Program (RTP) scholarship. SLC is supported by Australian Antarctic Science Grant 4482. SK thanks the South African National Department of Environment Affairs through its funding of the South African National Biodiversity Institute’s Directorate for Biological Invasions and the DSI-NRF Centre of Excellence for Invasion Biology (CIB) at Stellenbosch University. HER was partly supported by the Natural Environment Research Council award number NE/R016429/1 as part of the UK-SCAPE programme delivering National Capability. AML received support from the National Science Foundation Macrosystems Biology Program (grant numbers 1241932, 1638702) and grant EVA4.0, No. CZ.02.1.01/0.0/0.0/16_019/0000803 financed by OP RDE.

